# CORONAVIRUS INFECTION ACCELERATES EXPERIMENTAL OSTEOPOROSIS THROUGH THE ACTIVATION OF OSTEOCLASTS

**DOI:** 10.64898/2026.10.28.762635

**Authors:** Felipe Emanuel Oliveira Rocha, Letícia Cristina Dias Lima, Bárbara Luísa Vieira da Silva, Viviane Lima Batista, Jenniffer Ramos Martins, Fernanda de Lima Tana, Felipe Rocha da Silva Santos, Joyce Elisa Heredia, Matheus Rodrigues Gonçalves, Danielle Cunha Teixeira, Maria Luiza Canalis, Guilherme de Paula Costa, Talita Martins, Diogo Maia Moreira dos Santos, Pedro Pires Goulart Guimarães, Clésia Cristina Nascentes, Eduardo Henrique Martins Nunes, Mauro Martins Teixeira, Vivian Vasconcelos Costa, Celso Martins Queiroz-Junior

## Abstract

Coronavirus infection is increasingly associated with systemic inflammation and extrapulmonary sequelae, including bone loss. However, the impact of coronavirus infection on hosts with pre-existing osteoporosis remains poorly understood. Here, we investigated the effects of infection with the murine coronavirus MHV-A59 on bone remodeling in a mouse model of ovariectomy-induced osteoporosis. Female C57BL/6 mice underwent ovariectomy (OVX) or sham surgery and, after establishment of the osteoporotic phenotype (14 days post-surgery), were intranasally infected with MHV-A59. Clinical, inflammatory, and femur parameters were evaluated throughout acute and chronic phases of infection using histopathological, micro-computed tomography and biochemical analyses. *In vitro* infection assays with bone-marrow derived macrophages (BMDM) and BMDM-differentiated osteoclasts from Sham and OVX mice were also performed. MHV-A59 induced comparable lung inflammation in Sham and OVX mice. In contrast, infection accelerated bone resorption in OVX mice during the acute phase (5 dpi), whereas in non-osteoporotic animals, infection induced significant bone loss at later stages (60 dpi). These alterations were associated with increased osteoclast differentiation and activity and with elevated levels of osteometabolic and inflammatory mediators, including RANKL, CCL2, CCL3, CCL5, and TGF-β1. *In vitro*, MHV-A59 infection enhanced the expression of osteoclast-associated markers, including *Rank*, *Traf6*, *Nfac1*, and cathepsin K, and inflammatory mediators by BMDM and osteoclasts. Importantly, these responses were significantly more pronounced in cells derived from OVX mice. Collectively, our findings demonstrate that coronavirus infection promotes bone resorption and accelerates early bone loss in the context of estrogen deficiency, highlighting enhanced osteoclast activity as a potential mechanism underlying this infection-induced phenotype.

**CLINICAL PERSPECTIVES:**

- Coronavirus infection can affect bone remodeling, but its impact on pre-existing osteoporosis remains unclear.
- MHV-A59 accelerated bone resorption in OVX mice and increased osteoclast-associated responses.
- These findings suggest that coronavirus infection may trigger and accelerate skeletal deterioration in healthy and estrogen-deficient individuals.

## INTRODUCTION

Infection with SARS-CoV-2, the etiological agent of COVID-19, emerged in late 2019 and rapidly evolved into a global pandemic, resulting in approximately 779 million cases and about 7 million deaths by September 2026 [1]. Although initially recognized as a respiratory disease, evidence accumulated throughout the pandemic has demonstrated that COVID-19 constitutes a multisystemic condition, capable of affecting various organs and tissues beyond the lungs, including the cardiovascular, renal, neurological, gastrointestinal, and musculoskeletal systems [2–5]. Recent evidence indicates that in some individuals, these extra-pulmonary symptoms may develop at least three months and, in some cases, years after the acute phase of COVID-19, which is called long COVID [6]. Epidemiological studies demonstrate that long COVID occurs more frequently in women [7] and these late manifestations also involve multiple systems [8–13].

Among the already suggested effects of long COVID are clinical and experimental evidence for bone alterations. Patients with COVID-19 present reduced serum levels of osteocalcin, increased sclerostin and DKK-1, decreased bone mineral density, and a probable higher incidence of osteoporosis after infection [14–17]. In parallel, experimental models have demonstrated that coronavirus infection induces bone loss associated with an increase in the number and activity of osteoclasts, elevated RANKL expression, and activation of TNF- and IL-1β-dependent inflammatory pathways [18–21]. In fact, murine coronavirus infects macrophages and induces osteoclast differentiation independently of RANKL [20]. Meanwhile, although SARS-CoV-2 induces an abortive infection phenotype in human macrophages, it can directly stimulate osteoclastic differentiation, and also compromise osteoblastic function, favoring an imbalance in bone remodeling towards resorption [22–23].

Despite these advances, the effects of coronavirus infection on individuals with pre-existing conditions of bone fragility, such as postmenopausal osteoporosis, remain insufficiently investigated. This gap is particularly relevant considering that postmenopausal women constitute a population highly susceptible to osteoporosis resulting from estrogen deficiency [24–25]. The incidence of fragility fractures increases progressively with age, becoming markedly higher from age 70 in women and age 80 in men [26]. Furthermore, older adults also constitute one of the main risk groups for COVID-19 and long COVID [27]. Indeed, a recent cross-sectional study reported that SARS-CoV-2 reinfection was independently associated with an increased risk of osteoporosis in individuals over 55 years of age [28]. Nevertheless, further evidence is still required to clearly support the hypothesis that coronavirus infection exacerbates osteoporosis.

Therefore, in the present study we aimed to investigate the effects of murine coronavirus infection on bone tissue in ovariectomy-induced osteoporotic mice. Our results demonstrated that coronavirus infection accelerates experimental osteoporosis through the activation of osteoclasts, and amplification of inflammatory pathways associated with bone resorption.

## METHODS

### Animals

All the experimental procedures performed in this study complied with the recommendations of CONCEA (National Council for the Control of Animal Experimentation) and were approved by the local Ethics Committee on the Use of Animals of UFMG (process n° 057/2023). Female C57BL/6 mice aged 5 weeks-old (n=5 per group) were kept in individual ventilated cages in the facilities of the Animal Facility of the Department of Biochemistry and Immunology of the Institute of Biological Sciences of UFMG, with a 12/12 hour light/dark cycle, at room temperature (23-25 °C), without restrictions on food and water.

### Ovariectomy

To study the effects of coronavirus infection on osteoporosis, an experimental model of osteoporosis induced by bilateral ovariectomy (OVX) in mice was used [29]. Briefly, the mice were anesthetized intraperitoneally with ketamine (80 mg/kg) and xylazine (15 mg/kg), followed by surgical removal of the ovaries. The animals in the control group, called Sham, also underwent a similar surgery procedure, but without ovary removal. To characterize the osteoporotic phenotype, mice were weighed throughout the experiment to assess weight gain or loss. After euthanasia, the uterus of the animals was collected and weighed (mg) in order to compare the values obtained between the experimental groups to determine if uterine atrophy occurred.

### Cells and viruses

Fibroblast cells from *Mus musculus* mice [L929 cell line; CCL-1™ – American Type Culture Collection (ATCC)] were cultured in a controlled atmosphere (37 °C, 5% CO[) in Dulbecco’s Modified Eagle Medium (DMEM; Cultilab), supplemented with 7% fetal bovine serum (FBS; Gibco), 100 U/mL penicillin, and 100 µg/mL streptomycin. This cell line was used for viral propagation and titration. The coronavirus A59 strain of murine hepatitis virus (MHV-A59), acquired from ATCC (VR-764), after propagation, was kept in a freezer at −80°C until use.

### Coronavirus infection

Mice were infected with coronavirus in a biological safety cabinet 14 days after the osteoporosis induction by ovariectomy. To that end, mice were anesthetized intraperitoneally with ketamine (80 mg/kg) and xylazine (15 mg/kg), and received an intranasal inoculation of 30 μL of the betacoronavirus MHV-A59 (3×10[ PFU) in sterile saline solution (Samtec Biotecnologia), while only saline was instilled in the simulated controls [30]. The weight of the mice was measured every day after infection (dpi) to assess body mass. Euthanasia occurred at 2, 5, and 60 dpi for collection of blood, plasma, lung, uterus, and femur.

### Euthanasia, sample collection and hematological parameters

Mice were anesthetized intraperitoneally [ketamine (80 mg/kg) and xylazine (15 mg/kg)], and blood samples were collected from the abdominal vena cava. Subsequently, cervical displacement was performed, and the organs collected for histological and micro-CT analyses were fixed in 10% v/v formalin (ACS Científica) (pH 7.4). Samples intended for ELISA and PCR were flash-frozen in liquid nitrogen and stored at −80°C. From the collected blood, total and differential leukocyte counts were performed using a Celltac MEK-6500K veterinary hemocytometer (Nihon Kohden). Blood samples were also centrifuged at 5.000 rpm for 10 minutes at 4°C, to collect the plasma for enzymatic immunoassays.

### Histological analysis

The tissues were fixed in 10% v/v buffered formalin (pH 7.4) for 48 hours, dehydrated in graded ethanol, and embedded in paraffin (VETEC). After fixation, the femurs underwent decalcification in 14% EDTA for 30 days before further tissue processing. Serial 5 μm thick sections were made and stained with hematoxylin and eosin (H&E), Masson’s trichrome, or tartrate-resistant acid phosphatase (TRAP) assay. Lung slides were stained with H&E and examined for inflammation-mediated injury using a scoring system encompassing (I) airway damage (up to 4 points), (II) vascular inflammation (up to 4 points), (III) parenchymal alterations (up to 5 points), and (IV) neutrophil infiltration (up to 5 points), as described by Queiroz-Junior *et al*. [20]. Bone slides stained with Masson’s trichrome were photographed in the distal femur region, near the epiphyseal cartilage, and evaluated for the percentage of trabeculae relative to the total area assessed and for osteoblast and osteocyte counts, using ImageJ software (National Institutes of Health). Osteoblasts were identified by their position within the bone trabeculae and by their morphological characteristics, while osteocytes were identified in lacunae of the mineralized bone matrix. The cells were counted and the data expressed as the number of cells per perimeter of bone trabeculae. The number of cells was normalized by the evaluated bone perimeter [Cells/B.Pm (#/mm) -magnification: 400×]. The TRAP assay (Tartrate-resistant acid phosphatase assay) was performed on femur samples and also on macrophage and osteoclast cell cultures. TRAP activity was evaluated using histochemical staining to identify osteoclasts. The reaction used a reaction buffer containing 0.05 M sodium acetate (Synth), 0.05 M sodium tartrate (Synth), and 0.1% acetic acid (Synth) (pH 4.7-5.0) with a staining solution with 1% (v/v) N,N-dimethylformamide (Synth), 0.5 mg/mL Fast Red Violet LB Salt (Sigma-Aldrich, Cat# F3381) and 0.1 mg/mL Naphthol AS-MX phosphate (Sigma-Aldrich, Cat# N4875). Osteoclasts were identified as TRAP-positive cells adjacent to bone trabeculae next to the distal metaphyseal region of the femur. The number of cells was expressed per bone trabecular perimeter evaluated in a total of 10 microscopic fields ([Cells/B.Pm (#/mm) -magnification: 400×]. For *in vitro* data, cells were photographed (magnification: 100×) to count TRAP-positive multinucleated cells (two or more nuclei) using ImageJ software.

### Micro-CT analysis

The femurs were fixed in 10% v/v formalin (pH 7.4) for 48 hours, washed with distilled water, and scanned using a compact desktop micro-CT scanner (SkyScan 1174, Bruker micro-CT) with a source voltage of 50 kV, a source current of 800 μA, and a pixel size of 8.05 μm. A 0.5 mm aluminum filter was used. The samples were fixed on a stage that rotated 180°, with images acquired every 0.7°. The acquired shadow projections (16-bit TIFF format) were subsequently reconstructed into 2D slices using the NRecon software interface (Skyscan, Bruker micro-CT). The CT-Analyser software (CTAn; Skyscan, Bruker Micro-CT) was used for 3D analyses, and CTVox (Skyscan, Bruker Micro-CT) was applied for 3D bone volume observation. The bone mineral density (BMD) was calibrated in CT-An against the attenuation coefficient, using a pair of 2 mm calcium hydroxyapatite phantom rods of known concentration (0.25 and 0.75 g/cm³ CaHA) scanned and reconstructed under the same settings as the specimens. For the microarchitecture analysis of the distal femurs, the chosen region of interest of the metaphyseal trabecular bone, containing 41 slices after the most proximal portion of the epiphyseal line, was evaluated in different parameters, including trabecular BMD (Tb.BMD), bone volume/total volume (BV/TV), number of bone trabeculae (Tb.N), trabecular thickness (Tb.Th), spacing between bone trabeculae (Tb.Sp), and percentage of porosity. Analysis of the cortical bone was conducted in the mid-diaphyseal region of the femur, delimited by the proximal and distal epiphyseal plates. Starting from the midpoint of the diaphysis, 100 consecutive sections were evaluated, 50 in the proximal direction and 50 in the distal direction, in which the following morphometric parameters were quantified: bone area (B.Ar), total cross-sectional area (T.Ar), bone area fraction (B.Ar/T.Ar), bone perimeter (B.Pm) total perimeter (T.Pm), and cortical thickness (Cs.Th).

### Immunohistochemistry analysis

Femur sections were processed for immunohistochemistry evaluation. Antigen retrieval was performed with 0.1 M citrate buffer solution (pH 6.0) for 20 minutes at 96°C. Endogenous peroxidase activity was blocked in 3% hydrogen peroxide solution, and nonspecific blocking was performed using Protein Block (Abcam, Cat# ab64226). The sections were then incubated with rabbit anti-mouse myeloperoxidase primary antibody (1:200; ThermoFisher Scientific; Cat# PA5-16672) or rabbit anti-mouse CD68 antibody (1:500; R&D System; Cat# MAB101141) and incubated at 4°C overnight. Next, the sections were incubated with the secondary antibody (VECTASTAIN Elite ABC-HRP Kit, Vector Laboratories, Cat# PK-6100) for 30 minutes, stained with 3,3′-Diaminobenzidine (DAB) chromogenic solution (Sigma-Aldrich, Cat# D7304), and counterstained with hematoxylin. For cell counting using ImageJ software, ten microscopic fields were photographed in the distal femur region, near the epiphyseal cartilage (magnification: 400×). Data were expressed as the average number of cells per mm² [Cells/T.A (#/mm^2^)].

### ELISA analysis

The quantification of mediators associated with bone metabolism, including RANKL (Cat# MTR00) and OPG (Cat# MOP00), as well as inflammatory cytokines, including IL-1β (Cat# MLB00C), IL-6 (Cat# M6000B), TNF-α (Cat# MTA00B), and TGF-β1 (Cat# DY1679), and chemokines, including CCL2 (Cat# MJE00B), CCL3 (Cat# MMA00), CCL5 (Cat# MMR00), and CXCL1 (Cat# MKC00B), was performed by Enzyme-Linked Immunosorbent Assay (ELISA) in both *in vivo* and *in vitro* experiments. All analytes were quantified using commercially available mouse-specific ELISA kits from R&D Systems. For *in vivo* analyses, lung and femur samples were homogenized using a TissueLyser LT (QIAGEN) in cytokine extraction buffer [0.4 M NaCl, 0.05% (v/v) Tween 20, 0.5% (w/v) BSA, 0.1 mM PMSF, 13 µM benzethonium chloride, 1 mM EDTA, 0.002% (v/v) aprotinin], containing protease inhibitor. The homogenates were centrifuged at 10,000 rpm for 10 min at 4 °C, and the resulting supernatants were collected for analysis. ELISA assays were performed according to the manufacturer’s instructions for each R&D Systems kit. For each analyte, mediator concentrations were determined by interpolating the absorbance values measured at 490 nm against the corresponding standard curve generated using the standards provided in the respective ELISA kit. Results were expressed as pg/mL for cell culture supernatants or as pg/100 mg of tissue for tissue samples.

### Culture of bone marrow-derived macrophages and osteoclast differentiation

Bone marrow-derived macrophages (BMDM) and BMDM-differentiated osteoclasts were cultured as described by Queiroz-Junior *et al*. [31]. Femurs from ovariectomized (OVX) and non-ovariectomized (Sham) mice (n=3/group) were collected and the obtained bone marrow cells were cultured in T-175 culture flasks (SARSTEDT) in DMEM 10% FBS supplemented with M-CSF (100ng/mL; Peprotech, Cat# 315-02) to induce the differentiation of myeloid precursors into BMDM. After 7 days, the cells were seeded at a density of 1 × 10[ cells/well in 96-well plates and 1 × 10[ cells/well in 24-well plates. For osteoclast differentiation, macrophages were stimulated with 50 ng/mL of receptor activator of nuclear factor κB ligand (RANKL) (R&D Systems, Cat#462-TEC) every two days for three times. For viral infection, macrophages and osteoclasts were exposed to MHV-A59 at a ratio of 1 viral particle per 100 cells [multiplicity of infection (MOI) of 0.01], in DMEM supplemented with 2% FBS. The plates were incubated for 1h in an incubator at 37 °C (5% CO[), with agitation every 15 min. Then, the viral inoculum was removed, the wells were washed with DMEM, and DMEM containing 10% FBS, with or without RANKL, was added in a final volume of 100 µL (96-well plate) or 500 µL (24-well plate) per well. The cells were incubated for different periods (12, 24 and 48h), the supernatant was collected and stored at −80 °C, and cell viability analyses, TRAP assay, PCR and quantification of viral titers and inflammatory mediators were performed.

### MTT cell viability assay

3-(4,5-dimethyl-2-thiazolyl)-2,5-diphenyl-2H-tetrazolium bromide (MTT, Invitrogen, Cat# M6494) was used to determine the viability of macrophages and osteoclasts cultured from the bone marrow of the femur of ovariectomized and non-ovariectomized mice infected with MHV-A59. Briefly, 100 μL of MTT solution (1 mg/mL final concentration) prepared in DMEM supplemented with 10% FBS was added to each well, and the plate was incubated at 37°C (5% CO_2_) for 1.5 hours. Then, the MTT solution was removed, and 100 μL of dimethyl sulfoxide (DMSO, ACS Científica) was added to solubilize the formazan crystals. The absorbance of the samples was obtained using a spectrophotometer (Biotek Epoch™) at 595 nm, and the results were expressed as a percentage of viable cells in relation to the control group.

### Viral titer

L929 cells were seeded in 24-well plates at a concentration of 1 x 10[ cells per well. After 24 hours, 100 μL/well of supernatant collected from macrophage and osteoclast cultures was added to the cells. The plates were incubated at 37°C (5% CO[) and shaken every 15 minutes for 1 hour. Subsequently, the supernatant was carefully removed, the wells were washed with 0% DMEM, and 1 mL/well of DMEM containing 0.8% carboxymethylcellulose (Synth), 2% FBS, and 1% penicillin/streptomycin (Sigma-Aldrich) was added. The plates were kept in the incubator for 3 days, after which they were fixed with 1 mL/well of 10% neutral buffered formalin for 1 hour and stained with 0.1% crystal violet (Synth). Virus titers were determined as PFU/100 µL of supernatant.

### RNA extraction and quantitative real-time PCR

Viral RNA from femur tissue was extracted using the QIAamp® Viral RNA Extraction Kit (QIAGEN, Cat# 52904), while total RNA from cultured macrophages and osteoclasts was extracted using TRIzol (ThermoFisher Scientific, Cat# 15596026) according to the manufacturer’s protocol. The quantity and purity of the extracted RNA were measured using a NanoDrop spectrophotometer (ND-1000; ThermoFisher Scientific). cDNA was synthesized from 500 ng of total RNA using the iScript^TM^ gDNA Clear cDNA Synthesis Kit (Bio-Rad, Cat# 1725035) according to the manufacturer’s instructions. Quantitative real-time PCR was performed using Fast SYBR^TM^ Green Master Mix (Applied Biosystems, Cat# 4385616) in a final volume of 10 μl, containing 1 μl of diluted cDNA (1:10). The primers, synthesized by Integrated DNA Technologies (IDT), were used at a final concentration of 5 nM, and the sequences employed for the detection of the target genes are described in **Table 1**. All reactions were performed in duplicate in a 7500 Fast thermocycler (Applied Biosystems), in which the amplification protocol consisted of one cycle at 95 °C for 20 min, followed by 39 cycles at 95 °C for 30 s and 60 °C for 1 min. The relative expression of the genes was determined by the 2-ΔΔCt method, and the data were normalized by the expression levels of the constitutive gene *Gapdh*.

**Table 1.**
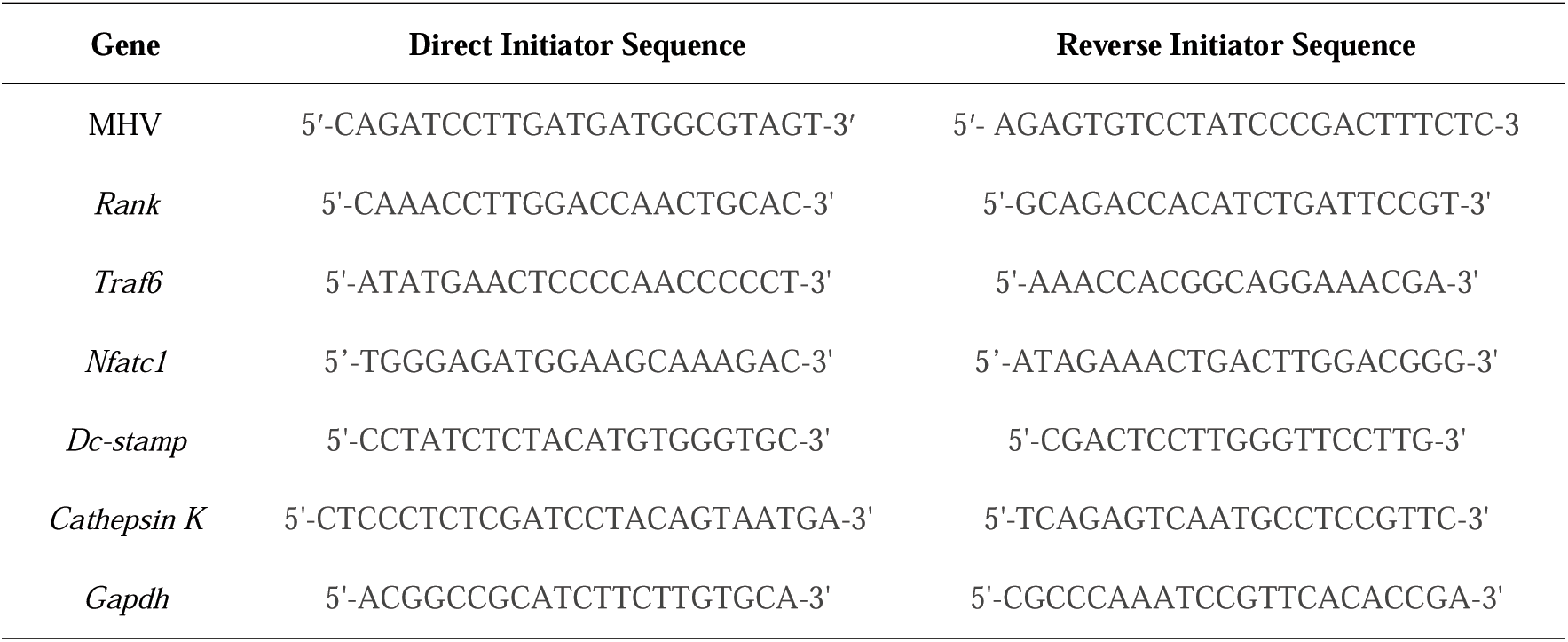
Primer Sequences.

### Transmission electron microscopy

BMDMs from Sham and OVX mice, previously differentiated as described, were seeded into T75 cell culture flasks (SARSTEDT) at a concentration of 8 × 10[ cells per flask and incubated in a 5% CO[ atmosphere at 37°C. After 24 hours, the cells were infected with MHV-A59 at a MOI of 0.01; 24 hours post-infection, they were fixed in 2.5% glutaraldehyde in 0.1 M phosphate buffer for 2 hours at room temperature. The cells were post-fixed with 2% osmium tetroxide and embedded in Epon resin. Ultrathin sections were then analyzed by TEM (Spirit BioTwin FEI, 120 kV) at the Microscopy Center of UFMG, Brazil.

### Statistical analysis

Statistical analyses were performed using GraphPad Prism software version 10.3.0 for Windows (San Diego, California, USA). Quantitative data were expressed as mean ± standard error of the mean (SEM). Initially, statistical analyses were performed to assess whether the obtained data had a normal distribution using the Shapiro-Wilk test. Differences between means were evaluated using the unpaired *Student’s t-test* or one-way ANOVA followed by *Tukey’s* multiple comparisons test when the data presented a normal distribution (parametric). Data that presented a non-normal distribution were evaluated using the non-parametric *Mann-Whitney* test followed by *Dunn’s* multiple comparisons test. Differences with *p* < 0.05 were considered statistically significant.

## RESULTS

### MHV-A59 INFECTION INDUCES BONE RESORPTION AND ACCELERATES OVARIECTOMY-INDUCED OSTEOPOROSIS THROUGH OSTEOCLAST ACTIVATION

The first step in this study was to investigate the systemic and lung phenotype of coronavirus infection in OVX mice (**Sup Fig 1A**). Ovariectomy (OVX) itself increased the body mass (**Sup Fig 1B**), induced leukocytosis (**Sup Fig 1D**) and uterine atrophy (**Sup Fig 1F**), when compared to non-ovariectomized animals (Sham). MHV-A59 infection induced a similar phenotype in both Sham and OVX groups, with mild weight loss (**Sup Fig 1C**) and significant leukopenia, but only the Sham-infected group showed increased granulocyte/lymphocyte ratio (**Sup Fig 1D-E**). When evaluating the pulmonary changes caused by the infection, similar mild to moderate lung damage occurred in the acute phase of infection and resolved at 60 dpi, in both Sham and OVX mice (**Sup Fig 1G**). Both groups presented hyperplasia of the alveolar walls associated with the infiltration of mononuclear and polymorphonuclear leukocytes, especially next to the perivascular and peribronchiolar regions (**Sup Fig 1H**). This is in line with previous published data [30]. When evaluating lung mediators responsible for such cell recruitment, a more robust chemokine response at 2 dpi was seen in Sham-infected mice (i.e. CXCL-1 (**Sup Fig 1I**), CCL2 (**Sup Fig 1J**), CCL3 (**Sup Fig 1K**), while delayed CCL3 and CCL5 increase (**Sup Fig 1K-L**) was seen in OVX-infected mice. Therefore, in general, the lung alterations induced by coronavirus infection were similar between Sham and OVX mice. Then, we investigated how systemic inflammation from MHV-A59 infection could contribute to bone changes induced by hormonal deficiency resulting from ovariectomy.

In line with literature evidence, OVX mice presented progressive reduction in femoral trabeculae, regardless of infection [32–33] (**Fig 1A**). Additionally, the mild coronavirus infection in Sham mice, also induced significant bone resorption detected at 60 dpi. Noteworthy, the coronavirus infection in OVX mice triggered an acceleration in bone loss, which was seen as early as at 5 dpi (**Fig 1A and 1F**). Nevertheless, in the chronic phase (60 dpi), OVX + MHV-A59 did not promote additional trabecular changes (**Fig 1A**). Along with the bone resorptive phenotype, a significant increase in the number of osteoclasts (TRAP[ cells) was observed in OVX animals at 2 and 5 dpi, being even higher in the OVX-infected group at 2dpi (**Fig 1B and 1G**). At 5 dpi, the infection itself in Sham mice also triggered an increase in the number of osteoclasts, predicting a bone resorptive environment, which was detected lately (60 dpi). Quantification of osteoblast and osteocyte populations demonstrated that, in the acute phase of the disease, neither ovariectomy nor infection promoted significant changes in the number of these cells (**Fig 1C-D and 1H**). At 60 dpi, osteocyte numbers decreased in the ovariectomized groups, regardless of infection (**Fig 1D**). All these bone changes were associated with low MHV-A59 detection in the femurs of infected animals at 5 dpi (**Fig 1E**). Together, these results show that coronavirus infection acts as an accelerator of bone loss rather than a determinant of its final severity in the context of ovariectomy-induced osteoporosis.

**Figure 1.**
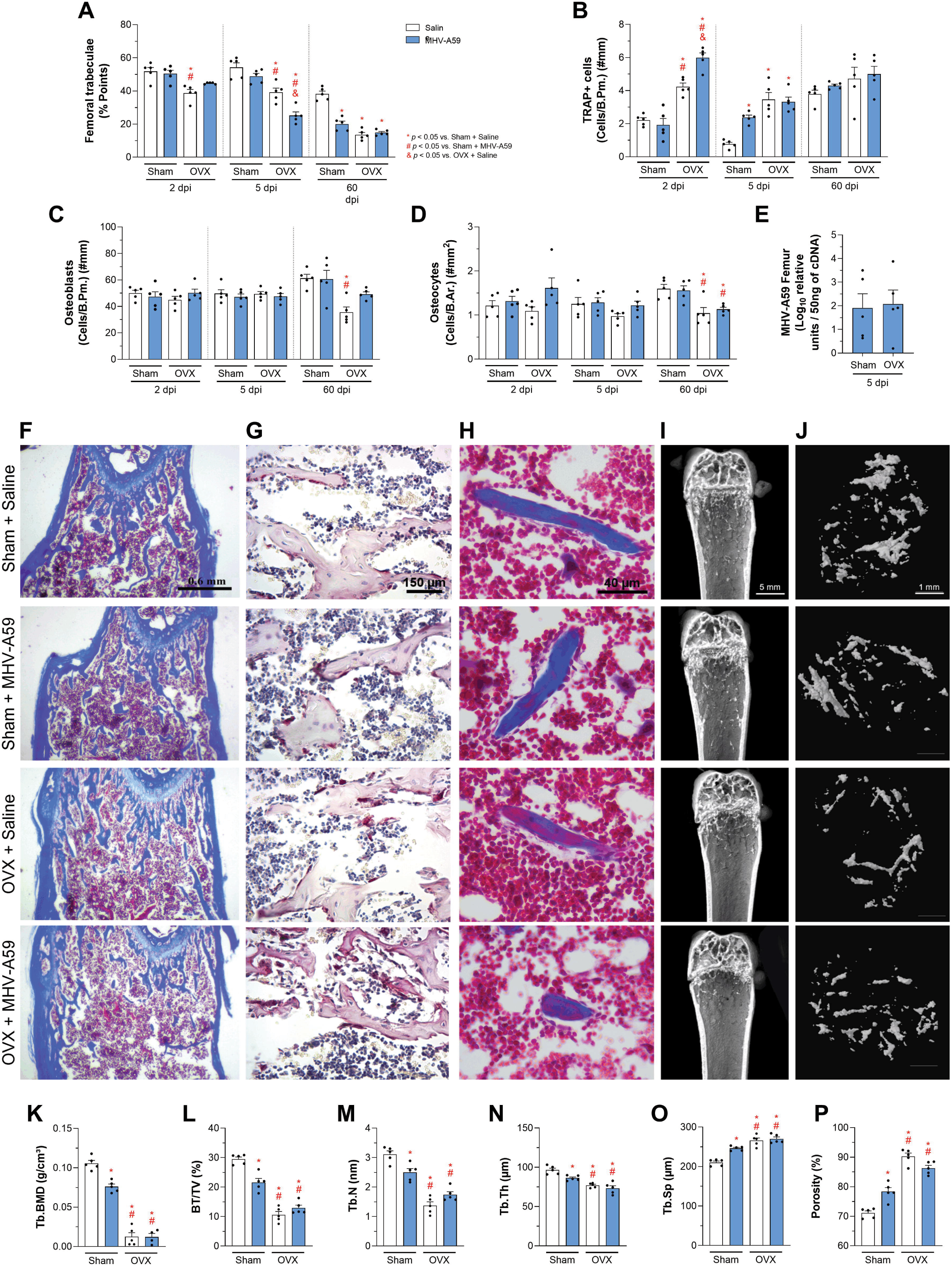
Effects of ovariectomy and MHV-A59 infection on histomorphometry, bone cellularity, viral load and trabecular microarchitecture of the femur. (A) Percentage of femoral trabeculae evaluated at 2, 5, and 60 days post-infection (dpi). (B-D) Quantification of bone lineage cells: osteoclasts (TRAP[) (B), osteoblasts (C), and osteocytes (D). Viral load in the femur 5 dpi (E). Representative images of histological sections of the femur stained with Masson’s Trichrome (F), TRAP for osteoclast identification (G), and osteoid/mineralized tissue staining (H), in Sham and OVX animals, control (saline) or infected with MHV-A59 5 dpi. Three-dimensional reconstructions by micro-CT of the distal femur (I) and trabecular region (J). Bone microarchitecture parameters obtained by micro-CT at 60 dpi, including trabecular bone mineral density (Tb.BMD) (K) bone volume/total volume (BV/TV) (L), number of bone trabeculae (Tb.N) (M), trabecular thickness (Tb.Th) (N), trabecular separation (Tb.Sp) (O), and percentage of bone porosity (P). White bars indicate control animals (saline) and blue bars indicate animals infected with MHV-A59. Points represent individual animal values (n = 5), and bars indicate mean ± standard error of the mean (SEM). Statistical differences are indicated by * (*p* < 0.05 vs. Sham + Saline), # (*p* < 0.05 vs. Sham + MHV-A59) and & (*p* < 0.05 vs. OVX + Saline).

To further investigate whether previous viral infection had left persistent effects on bone quality in OVX mice in the long term, we also assessed bone microarchitecture at 60 dpi by micro-CT (**Fig 1I and 1J**). The infection with MHV-A59 itself in Sham mice, as well as the OVX status, induced a significant reduction of trabecular bone mineral density (**Fig 1K**) and bone volume (BV/TV) (**Fig 1L**), a decrease in the number of trabeculae (**Fig 1M**), reduced trabecular thickness (**Fig 1N**), increased separation between trabeculae (**Fig 1O**), and greater bone porosity (**Fig 1P**), when compared to uninfected Sham animals. No differences were detected between non-infected and infected OVX mice. Additionally, estrogen deficiency and MHV-A59 infection did not promote any changes in cortical bone (**Sup Fig 2**). Therefore, despite inducing significant bone loss, coronavirus infection does not result in a sustained aggravation of the osteoporotic phenotype caused by ovariectomy.

### CORONAVIRUS INFECTION INCREASES OSTEOMETABOLIC AND INFLAMMATORY MEDIATORS AND ALTERS THE CELL POPULATIONS IN THE BONE MARROW

In order to investigate potential mechanisms underlying the accelerated bone loss observed during the acute phase of infection in OVX mice, we next evaluated the local expression of key mediators involved in bone remodeling. MHV infection induced a significant increase in femur RANKL levels in both Sham and OVX mice at 2 dpi compared with the Sham + Saline group (**Fig 2A**). OPG levels remained unchanged, while the RANKL/OPG ratio showed a biological increase at 5dpi that did not reach statistical significance **(Fig 2B-C)**. Additionally, infection with MHV-A59 triggered a significant increase in CCL2 levels at 2 dpi, which was followed by decrease in OVX and in infected mice at 5 dpi (**Fig 2D**). CCL3 and CCL5 levels significantly increased only in OVX-infected animals at 2 and 5 dpi, respectively (**Fig 2E and 2F**). When evaluating TGF-β1 levels, at 2 dpi, infection in Sham animals promoted a significant reduction in this cytokine, while in OVX animals, infection induced a significant increase in TGF-β1 at 5 dpi (**Fig 2G**). Additionally, the levels of other pro-inflammatory mediators, including IL-1β, IL-6, and CXCL-1, did not show significant variations among the groups (**Table 2**). Collectively, these results indicate that MHV infection elicits an inflammatory microenvironment in the femur, with infected OVX mice displaying a more robust chemokine response during the acute phase of infection.

**Figure 2.**
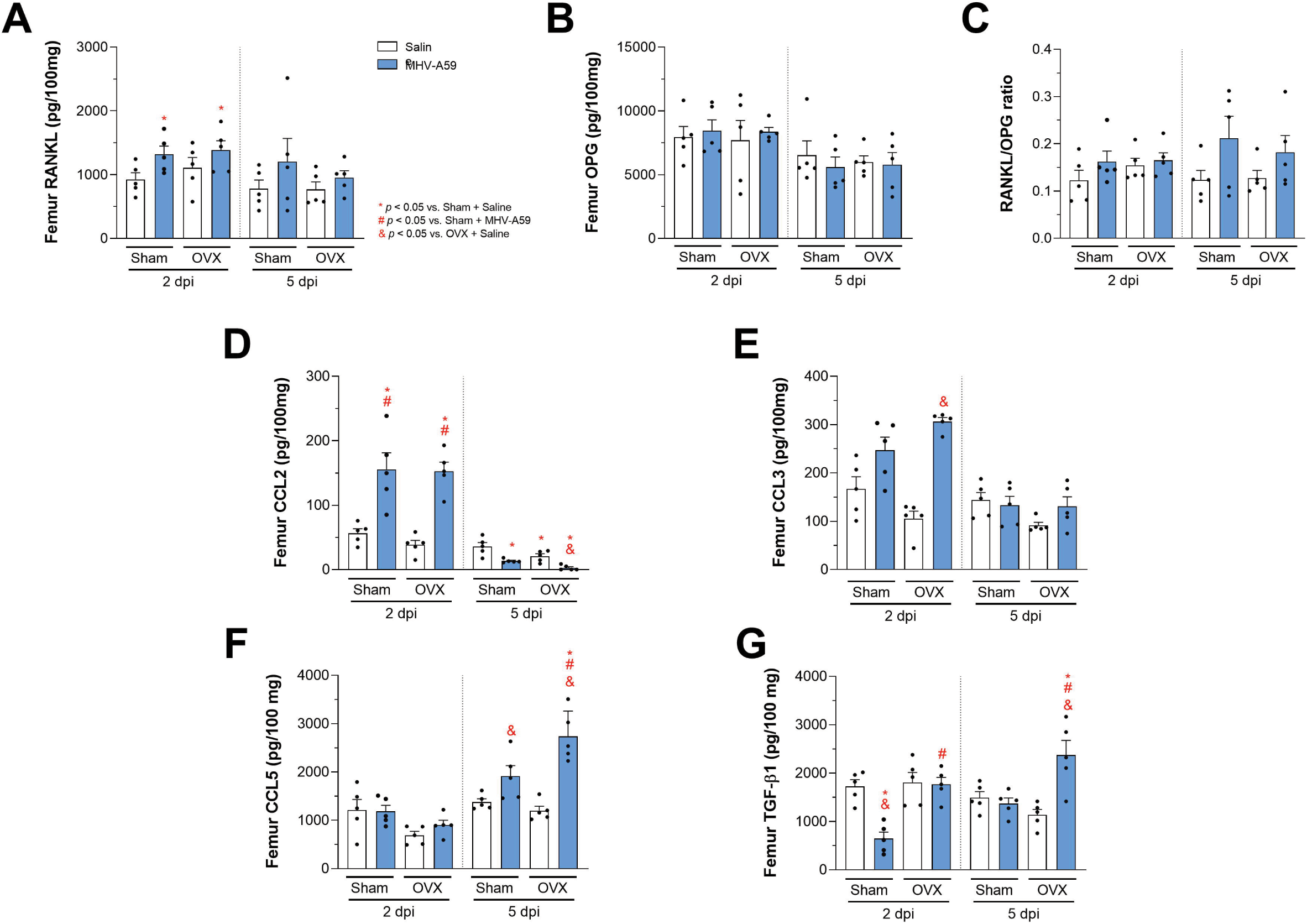
Profile of osteometabolic and inflammatory mediators in the femur of Sham and OVX mice after MHV-A59 infection. RANKL (A), OPG (B), RANKL/OPG ratio (C), CCL2 (D), CCL3 (E), CCL5 (F), and TGF-β (G) levels were quantified in the femur of Sham and OVX animals inoculated with saline or infected with MHV-A59 at 2 and 5 days post-infection (dpi). White bars indicate control animals (saline) and blue bars indicate animals infected with MHV-A59. Points represent values for individual animals (n = 5), and bars indicate mean ± standard error of the mean (SEM). Statistical differences are indicated by * (*p* < 0.05 vs. Sham + Saline), # (*p* < 0.05 vs. Sham + MHV-A59) and & (*p* < 0.05 vs. OVX + Saline).

**Table 2.**
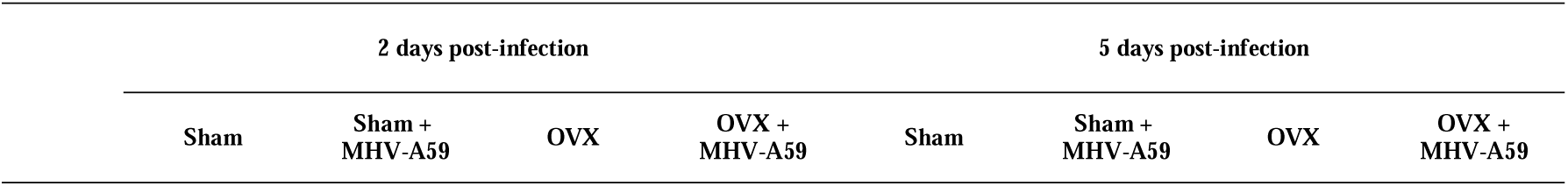
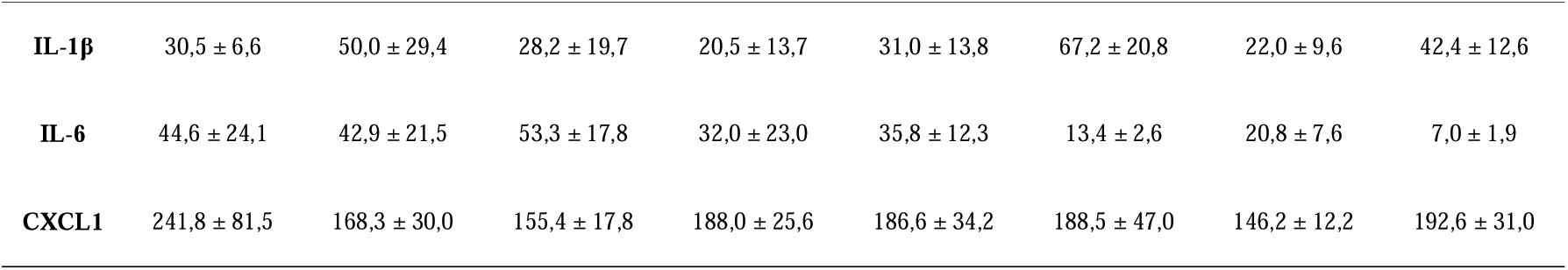
Pro-inflammatory mediator levels (pg/100 mg) in the femur of Sham and ovariectomized (OVX) mice at 2 and 5 days post-infection with MHV-A59.

The morphological and biochemical changes observed in the femur were also associated with alterations in bone marrow cell populations. Initially, MPO[ cells, which comprise predominantly neutrophils and granulocytic myeloid precursors in the bone marrow, were assessed. Infection significantly increased their numbers, with the OVX + MHV group exhibiting the highest values at 2 dpi (**Fig 3A-B**). Additionally, analysis of the CD68[ cell population, predominantly composed of monocytes, macrophages, and osteoclast precursors, revealed reduced cell numbers in both ovariectomized and infected animals, with the greatest reduction observed in OVX + MHV mice at 5 dpi (**Fig 3C-D**). These findings may reflect increased differentiation of monocyte/macrophage precursors into osteoclasts, along with infection-induced changes in monocytic populations.

**Figure 3.**
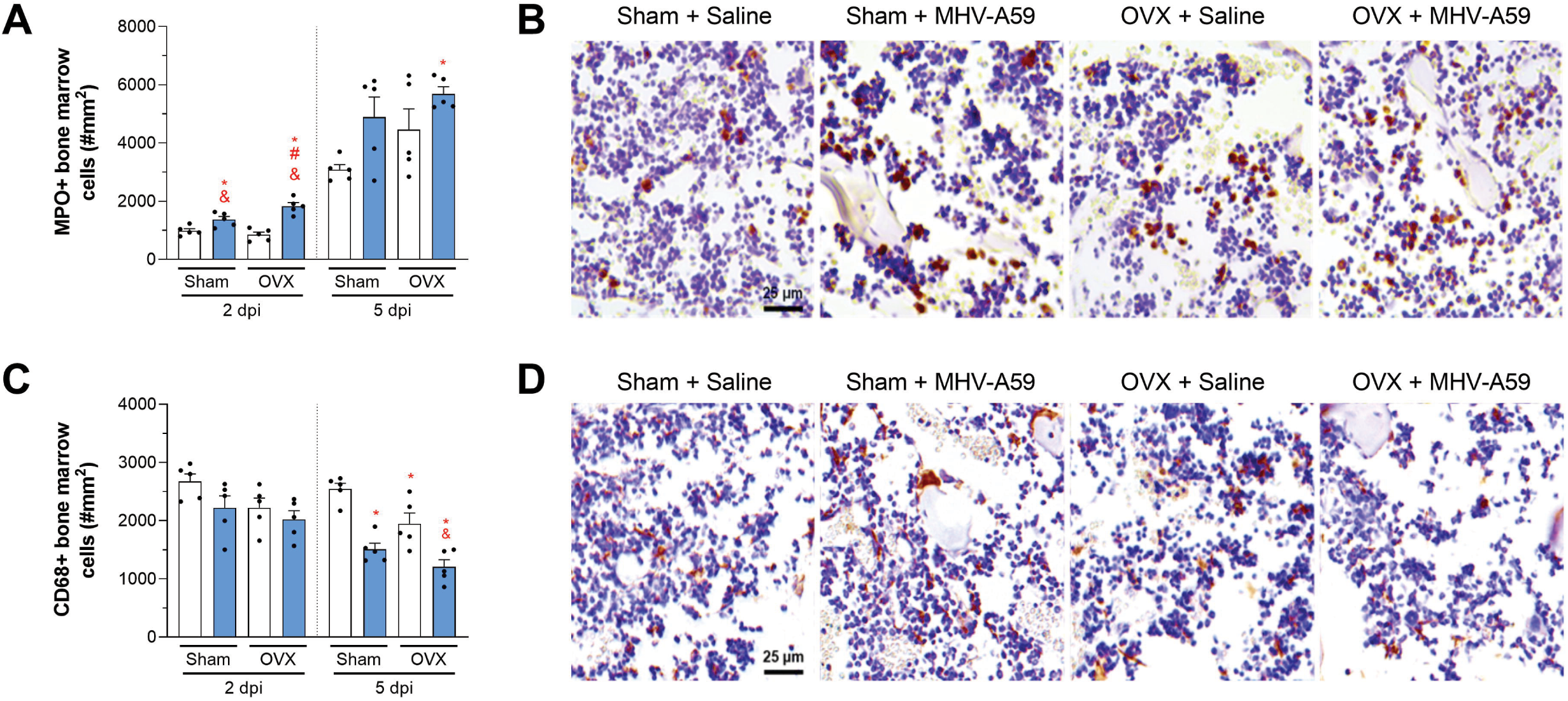
Modulation of MPOD and CD68D cells in the bone marrow of Sham and OVX mice after MHV-A59 infection. (A) Quantification of MPO[ cells in bone marrow at 2 and 5 days post-infection (dpi) in Sham and OVX animals infected with MHV-A59. (B) Representative immunohistochemistry images for MPO in bone marrow at 2 dpi. (C) Quantification of CD68[ cells in bone marrow at the same times and experimental groups. (D) Representative immunohistochemistry images for CD68 at 5 dpi. White bars indicate control animals (saline) and blue bars indicate animals infected with MHV-A59. Points represent individual animal values (n = 5), and bars indicate mean ± standard error of the mean (SEM). Statistical differences are indicated by * (*p* < 0.05 vs. Sham + Saline), # (*p* < 0.05 vs. Sham + MHV-A59) and & (*p* < 0.05 vs. OVX + Saline).

### MHV-A59 INFECTION PROMOTES THE DIFFERENTIATION OF MACROPHAGES FROM OVARIECTOMIZED MICE IN OSTEOCLASTS

Given the increased bone resorption observed *in vivo*, bone marrow-derived macrophages and RANKL-differentiated osteoclasts from Sham and OVX mice were infected with MHV-A59 (MOI 0.01), and TRAP[ cells, cell viability, and viral titers were evaluated (**Fig 4A**). Infection with MHV-A59 in macrophages induced an increase in the number of TRAP[ cells in cultures derived from OVX animals independent of RANKL (**Fig 4B**). Regarding RANKL-differentiated osteoclasts, the number of TRAP[ cells remained high in all groups (**Fig 4C**), with control OVX cells showing higher quantities compared to the control Sham group, as shown in **Fig 4D**, indicating that prior estrogen deficiency supports greater osteoclastogenic potential.

**Figure 4.**
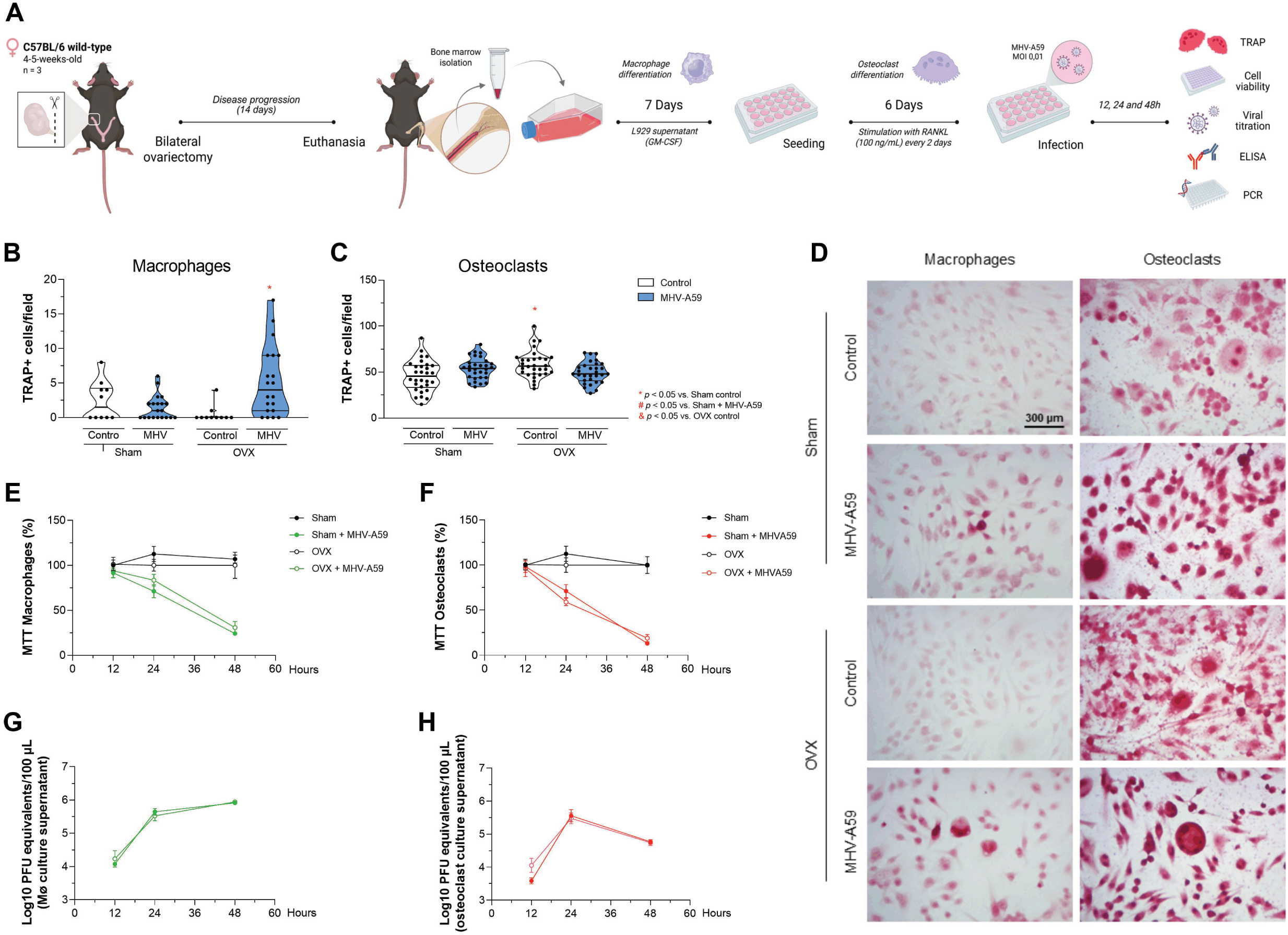
In vitro infection with MHV-A59 in macrophages and osteoclasts derived from the bone marrow of Sham and OVX mice. (A) Experimental design for isolation of bone marrow cells, in vitro differentiation into macrophages (L929 supernatant; GM-CSF) and osteoclasts (RANKL), followed by infection with MHV-A59. (B–C) Quantification of TRAP[ cells in macrophage and osteoclast cultures 24 hours after MHV-A59 infection. White bars indicate control (uninfected) cells and blue bars indicate MHV-A59 infected cells. Dots represent the average number of TRAP[ cells quantified in a total of 10 fields per sample. (D) Representative images of TRAP staining in macrophages and osteoclasts derived from Sham and OVX animals, under control conditions and after MHV-A59 infection. (E–F) Cell viability assay (MTT) in macrophages and osteoclasts, demonstrating the effect of infection over time. (G–H) Viral titer in the supernatant of macrophage and osteoclast cultures, indicating MHV-A59 replication in these cells. Black lines represent control (uninfected) cells, while green and red lines represent infected macrophages and osteoclasts, respectively. Data are presented as mean ± standard error of the mean (SEM). Statistical differences are indicated by * (*p* < 0.05 vs. Sham control), # (*p* < 0.05 vs. Sham + MHV-A59) and & (*p* < 0.05 vs. OVX control).

When evaluating cell viability and viral titers, MHV-A59 infection promoted a progressive and similar reduction in viability (**Fig 4E-F**) and increase in the viral titers in both macrophages and osteoclasts from sham or OVX mice over time (**Fig 4G-H**). Both cell types proved permissive to infection, sustaining the production of new MHV-A59 particles and therefore potentially acting as cellular reservoirs for the virus.

### PREVIOUS ESTROGEN DEFICIENCY AND MHV-A59 INFECTION INDUCE INTRACELLULAR CHANGES IN MACROPHAGES

To investigate potential ultrastructural changes underlying the enhanced osteoclastogenic potential of OVX-derived macrophages in response to MHV-A59 infection, these cells were analyzed by transmission electron microscopy. Qualitative analysis revealed specific ultrastructural changes in macrophages from Sham and OVX animals, as well as modifications associated with MHV-A59 infection. In the infected groups, regardless of hormonal status, macrophages exhibited irregular cell surfaces with numerous cytoplasmic projections (black arrows; **Fig 5A**), consistent with an activated cellular phenotype. Furthermore, cytoplasmic vesicles containing viral particles (green asterisks; **Fig 5D**) and viral particles associated with the nuclear membrane (red arrow; **Fig 5E**) were also observed, characteristics of the coronavirus replicative cycle.

**Figure 5.**
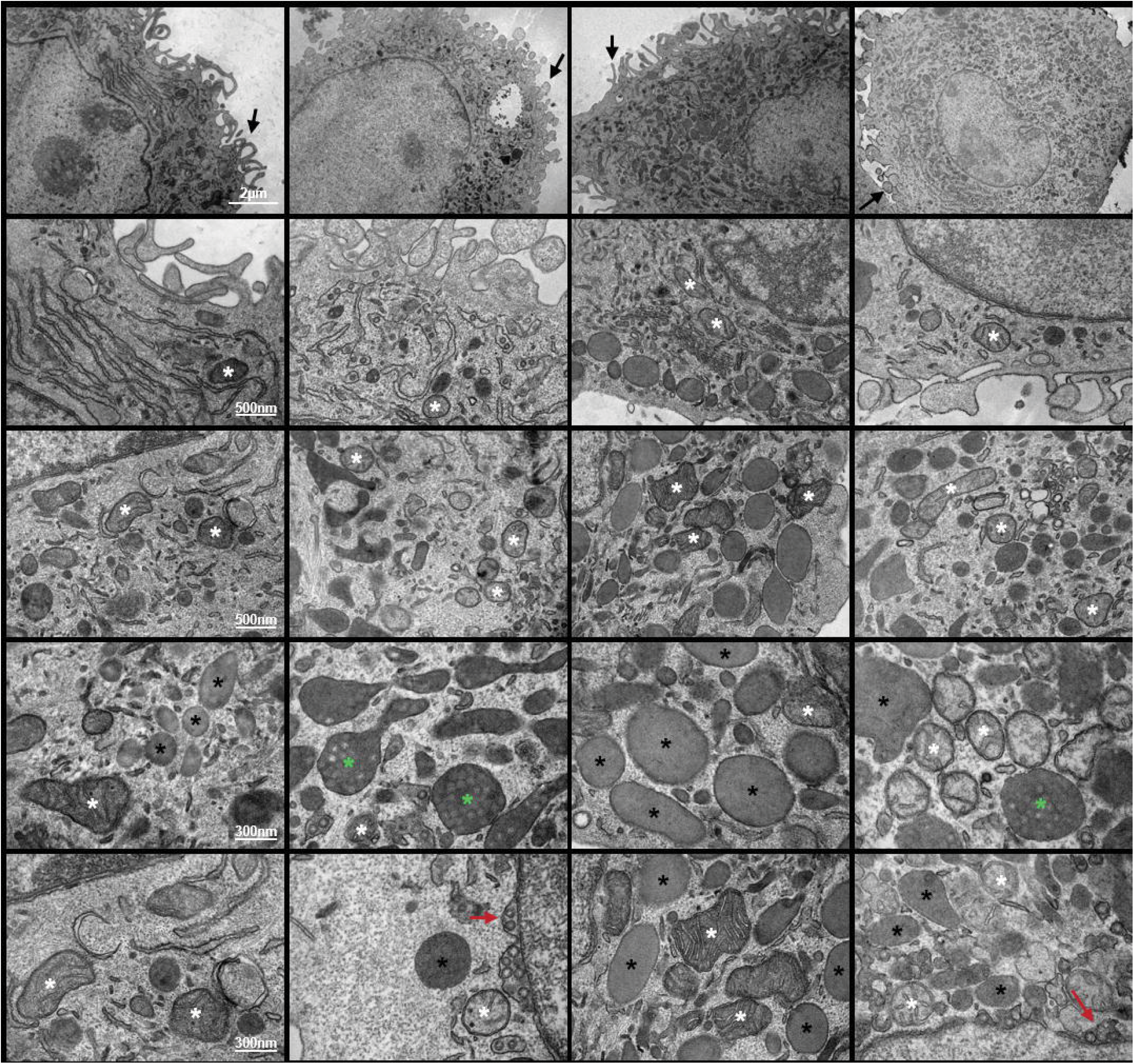
Transmission electron microscopy of bone marrow-derived macrophages from Sham and ovariectomized (OVX) mice, infected or not with MHV-A59. (A) Representative overview of macrophages showing alterations in the plasma membrane surface and the presence of numerous cytoplasmic projections (black arrows) in infected groups (B–C) Higher-magnification images highlighting the cytoplasmic organization and mitochondria (white asterisks), demonstrating increased mitochondrial abundance and morphological alterations in macrophages derived from OVX mice. (D) Lipid bodies (black asterisks), which were abundant in OVX groups, and cytoplasmic vesicles containing viral particles (green asterisks) observed in infected groups. (E) Viral particles associated with the nuclear membrane (red arrows). Scale bars: (A) 2 μm; (B-C) 500 nm; (D-E) 300 nm.

On the other hand, marked differences were evident between the cells derived from Sham and OVX groups. Macrophages from OVX animals exhibited prominent mitochondrial profiles (white asterisks; **Fig 5C**), including morphological changes suggestive of organelle remodeling, as well as conspicuous lipid droplet accumulation (black asterisks; **Fig 5D**). These features may reflect metabolic reprogramming that could be associated with previous estrogen deficiency. In contrast, macrophages from Sham animals showed mitochondria with preserved morphology and fewer apparent lipid inclusions, consistent with a less altered basal metabolic state.

Taken together, these findings suggest that previous estrogen deficiency promotes intrinsic ultrastructural and metabolic changes in macrophages, manifested by mitochondrial remodeling and lipid droplet accumulation, establishing a phenotype that is more susceptible to osteoclast differentiation. In contrast, MHV-A59 infection induces morphological changes typical of the viral replication cycle, without noticeably altering the differential ultrastructural pattern previously established by estrogen deficiency.

### INFECTION WITH MHV-A59 ALTERS THE EXPRESSION OF GENES ASSOCIATED WITH OSTEOCLAST DIFFERENTIATION AND ACTIVITY

The effects of infection were also studied in relation to the expression of genes associated with osteoclast differentiation in macrophage and osteoclast cultures derived from Sham and OVX mice. In macrophages, the infection induced a significant increase in the expression of *Rank*, *Traf6*, and *Nfatc1* (**Fig 6A-C**), independent of the cells’ previous hormonal status, while DC-STAMP expression did not differ among the groups (**Fig 6D**). Noteworthy, only macrophages derived from OVX mice showed an increase in Cathepsin K expression after MHV-A59 infection (**Fig 6E**).

**Figure 6.**
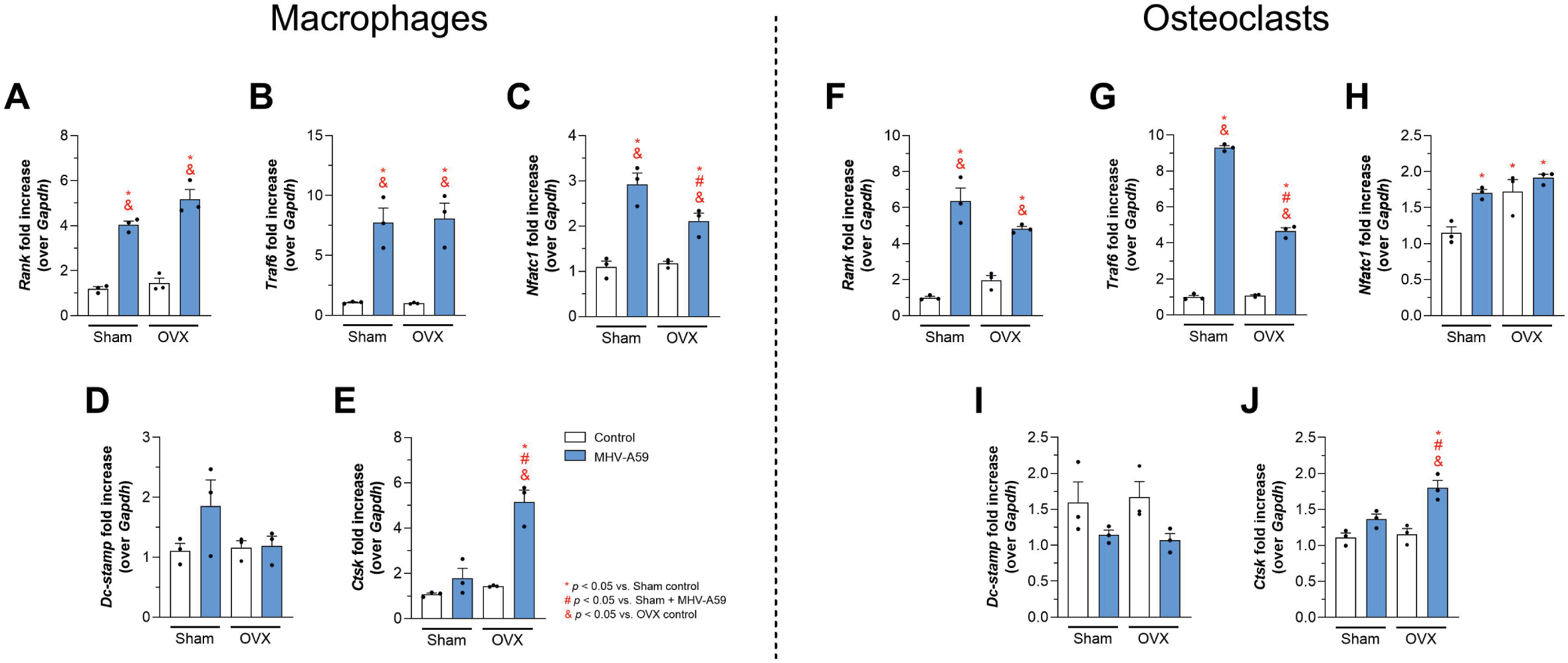
Gene expression of osteoclastogenic markers in macrophages and osteoclasts from bone marrow-derived Sham and OVX animals infected with MHV-A59 after 12 hours. Expression of *Rank* (A;F), *Traf6* (B;G), *Nfatc1* (C;H), *Dc-stamp* (D;I), *Cathepsin K* (*Ctsk*) (E;J), in macrophages and osteoclasts, normalized by *Gapdh* expression from control Sham cells according to cell line. White bars indicate control animals and blue bars indicate animals infected with MHV-A59. Points represent individual cell values (n = 3) and bars indicate mean ± standard error of the mean (SEM). Statistical differences are indicated by * (*p* < 0.05 vs. Sham control), # (*p* < 0.05 vs. Sham + MHV-A59) and & (*p* < 0.05 vs. OVX control).

In osteoclasts, the infection also induced increased expression of *Rank* and *Traf6*, (**Fig 6F-G**). While *Nfatc1* expression was intrinsically elevated in osteoclasts from OVX mice, MHV infection was sufficient to induce a similar increase in osteoclasts derived from Sham animals (**Fig 6H**). Ovariectomy, as well as MHV-A59, did not cause changes in DC-STAMP expression (**Fig 6I**). Regarding *Cathepsin K*, as expected, RANKL treatment significantly increased its expression when compared with macrophages (Sham macrophages: 1.11 ± 0.1 vs. Sham osteoclasts: 68.6 ± 13.9). When evaluating the effect of infection in osteoclasts themselves, MHV increased *Cathepsin K* expression only in osteoclasts derived from OVX mice (**Fig 6J**). Collectively, MHV infection enhanced the osteoclastogenic activity, both in macrophages and osteoclasts, particularly derived from OVX mice.

### MHV-A59 INFECTION INDUCES AN INCREASE IN INFLAMMATORY MEDIATORS IN MACROPHAGES AND OSTEOCLASTS FROM SHAM AND OVX ANIMALS

In macrophages, MHV-A59 infection induced a significant increase in IL-6 and TNF production in cells derived from both Sham and OVX groups (**Fig 7A-B**). Notably, infected OVX macrophages produced higher levels of IL-6 than infected Sham cells, indicating that previous estrogen deficiency amplifies the IL-6 response to infection. In contrast, neither infection nor ovariectomy altered CCL2 production (**Fig 7C**). Basal CCL3 levels were significantly increased in OVX macrophages compared to Sham cells, and infection further enhanced CCL3 production in both groups, with OVX macrophages exhibiting a stronger response when compared to infected Sham macrophages (**Fig 7D**). MHV-A59 infection also promoted a significant increase in CCL5 production, although this response was more pronounced in Sham macrophages than in OVX macrophages (**Fig 7E**).

**Figure 7.**
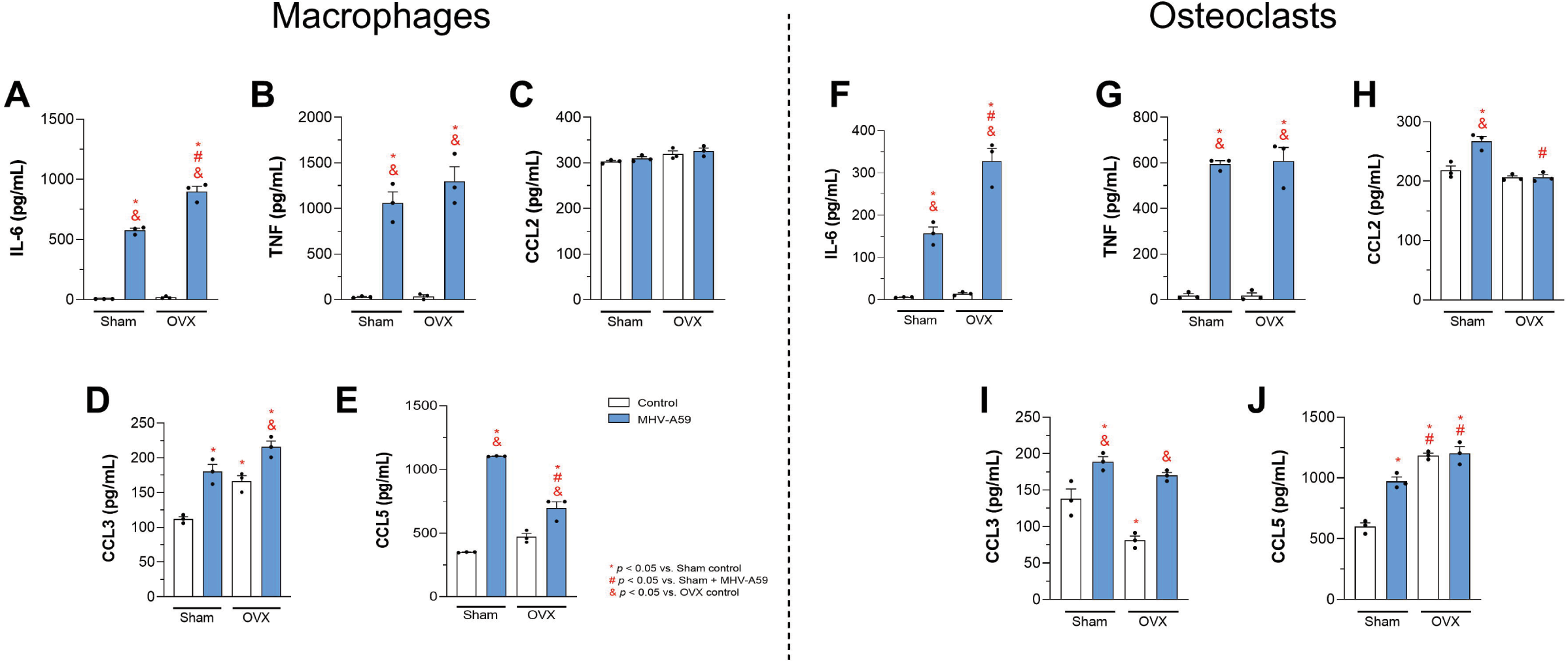
Secretion of cytokines and chemokines in the supernatant of macrophage and osteoclast cultures from bone marrow derivatives of Sham and OVX animals 48 hours after MHV-A59 infection. (A;F) IL-6, (B;G) TNF, (C;H) CCL2, (D;I) CCL3, and (E;J) CCL5 were quantified by ELISA in the supernatant of macrophages and osteoclasts. White bars indicate control animals and blue bars indicate animals infected with MHV-A59. Points represent individual cell values (n = 3) and bars indicate mean ± standard error of the mean (SEM). Statistical differences are indicated by * (*p* < 0.05 vs. Sham control), # (*p* < 0.05 vs. Sham + MHV-A59) and & (*p* < 0.05 vs. OVX control).

In osteoclasts, MHV-A59 infection also increased IL-6 and TNF production in cells derived from both Sham and OVX groups (**Fig 7F-G**). Similar to macrophages, infected OVX osteoclasts produced significantly more IL-6 than Sham-infected osteoclasts. Analysis of chemokine production revealed a distinct profile compared to macrophages. Infection increased CCL2 production in Sham-derived osteoclasts (**Fig 7H**). Basal CCL3 levels were reduced in OVX osteoclasts compared to Sham cells, while infection increased levels both in sham and OVX derived cells (**Fig 7I**). Finally, the previous OVX status and the infection increased CCL5 production by osteoclasts, in relation to Sham-derived osteoclasts (**Fig 7J**). Together, these findings demonstrate that previous estrogen deficiency partially favors the inflammatory and chemokine responses in macrophages and osteoclasts during MHV-A59 infection.

## DISCUSSION

Viral infections can affect bone tissue through inflammatory and osteoclastogenic pathways; however, although coronavirus infection has been associated with bone loss and increased osteoclastogenesis [18,20], its impact in individuals with pre-existing osteoporosis remains poorly understood. Here, we show that MHV-A59 lung infection promotes bone resorption and differentially affects bone remodeling according to estrogen status. Infection accelerated bone resorption in OVX mice during the acute phase of the lung disease, whereas infected and non-infected OVX mice exhibited a similar osteoporotic phenotype at 60 dpi. This response was associated with increased osteoclast numbers and activity, local induction of osteometabolic and inflammatory mediators, and enhanced osteoclastogenic responses in macrophages and osteoclasts derived from OVX mice. These findings indicate that estrogen deficiency increases susceptibility to an early infection-induced pro-resorptive response without further aggravating the long-term bone disease.

Bone alterations have emerged as a potential component of the long-term consequences of coronavirus infection. Clinical studies have reported reduced osteocalcin levels, increased sclerostin and DKK-1, decreased bone mineral density, and a potentially increased risk of osteoporosis following COVID-19 [14–17]. Experimental studies have provided mechanistic support for these observations, demonstrating that coronavirus infection can promote bone loss through osteoclast activation and inflammatory pathways [18–19,21]. Consistent with our previous findings showing that acute MHV-3 infection induces a TNF-dependent osteoporotic phenotype [20], the present study further demonstrates that coronavirus-induced bone resorption may be particularly relevant in the context of estrogen deficiency. This issue is clinically important because older women represent a population at increased risk of both osteoporosis and persistent post-COVID manifestations [7,24–25,27]. Notably, although infection accelerated bone loss in OVX mice during the acute phase, the observed phenotype of infected and non-infected OVX animals converged at 60 dpi, indicating that coronavirus infection primarily accelerates the progression of bone loss rather than determining its long-term severity. This interpretation is consistent with experimental evidence showing that coronavirus infection induces bone resorption through increased osteoclast activity [18–21] and with recent clinical evidence associating SARS-CoV-2 reinfection with an increased risk of osteoporosis in individuals over 55 years of age [28].

The observed bone phenotype was not related to worse lung disease in OVX mice. We have previously shown that MHV-A59 lung lesion and viral load are similar between Sham and OVX mice [30]. This is partially different from clinical data that showing that a high level of estradiol protects young female COVID-19 patients, while estrogen loss in older women may increase fatality rate of women. AMH and E2 estrogen receptors may protect against severe COVID-19, with E2 potentially exerting this effect through the modulation of immune and inflammatory cytokines [34–35]. In fact, here we have detected a delayed chemokine response to the virus after ovariectomy, and delayed expression of chemokines and cytokines can induce an inappropriate inflammatory response and lung disease during SARS-CoV infection [36]. Nevertheless, this chemokine alteration did not result in more severe pulmonary disease in OVX mice, possibly because MHV-A59 mimics a mild COVID-19 disease, inducing a transient inflammatory response with mild pulmonary disease in mice, consistent with other studies using such murine coronavirus model [30,37]. Thus, the differences observed in bone appear to be mainly related to the interaction between the infection-induced response and an osteoclastogenic microenvironment previously conditioned by estrogen deficiency.

The transient increased response of bone resorption induced by MHV-A59 infection in OVX mice was accompanied by an acute increase in femur RANKL, despite no significant changes in OPG or in the RANKL/OPG ratio. Thus, rather than indicating a global disruption of the RANKL/OPG axis, the increase in RANKL may have contributed to enhanced osteoclast precursor differentiation and activity within a microenvironment already predisposed to bone resorption, potentially reflecting the host response to viral entry into the bone marrow, even at the low levels detected. Indeed, numerous inflammatory cytokines and chemokines can promote osteoclast formation and bone degradation, being RANKL considered the final downstream effector cytokine that drives osteoclastogenesis and osteoclastic bone resorption [38]. The pro-resorptive response in OVX-infected mice was supported by increased CCL3 and CCL5 levels in the femur. CCL3 can enhance osteoclastogenesis and osteoclast survival, while also exerting negative effects on the osteoblastic lineage [39–45]. CCL5 also has pleiotropic functions and may promote myeloid cell recruitment, osteoclastogenesis, and the expression of matrix metalloproteinases [46–50]. These changes in the inflammatory and remodeling environment were accompanied by alterations in the bone marrow myeloid compartment, characterized by increased MPO[ and reduced CD68[ cell populations, concomitant with enhanced osteoclastogenesis, suggesting a cellular reorganization associated with the pro-resorptive response to infection. While MPO increase may be generally associated to a response with reactive oxygen species and pro-osteoclastogenesis [51–54], the reduction in the CD68[ population may reflect cell mobilization, differentiation, or infection-induced cell death [20]. However, these populations should not be interpreted as direct surrogates of osteoclastogenesis, since MPO is expressed across different stages of myeloid differentiation and CD68[ cells encompass multiple populations of the mononuclear phagocyte system [55–56]. The concomitant increase in TGF-β1 may also reflect the enhanced osteoclastic activity, given that this cytokine is abundantly stored in the bone matrix and released during bone resorption, while also participating in the regulation of RANKL expression and the coupling of bone remodeling [57–59]. Overall, these findings suggest that MHV-A59 infection transiently amplifies bone resorption in estrogen-deficient mice through coordinated changes in RANKL, inflammatory chemokines, TGF-β1, and the bone marrow myeloid compartment.

The *in vitro* findings provide a mechanistic link to the accelerated bone loss observed in OVX mice during the acute phase of MHV-A59 infection. Bone marrow-derived macrophages from OVX animals showed greater differentiation into TRAP[ osteoclasts following MHV-A59 stimulation in the absence of exogenous RANKL. Importantly, this difference was not associated with greater viral replication in OVX cells, indicating that previous estrogen deficiency enhances the osteoclastogenic response to infection rather than simply increasing cellular susceptibility to the virus. The RANKL-independent increase in osteoclast formation is consistent with previous evidence that coronavirus infection can directly promote osteoclastogenic programs in myeloid cells [20,22], while also extending these observations by showing that previous estrogen deficiency may amplify this response. Indeed, the ultrastructural analysis revealed that previous estrogen deficiency and viral infection exerted distinct effects on macrophage morphology. MHV-A59 induced membrane remodeling and cytoplasmic alterations consistent with the replication of coronaviruses [60–61], whereas OVX-derived macrophages exhibited increased mitochondrial content and lipid droplet accumulation regardless of infection. These features are consistent with the metabolic adaptations associated with macrophage differentiation into osteoclasts, which involves changes in mitochondrial activity and lipid metabolism and is modulated by estradiol [62–64]. Additionally, lipid droplets have been shown to favor the replicative cycle and immune response in monocytes after coronavirus infection [65], despite no difference in viral titers between sham and OVX-derived cells have been detected. Although these observations are based primarily on morphological and qualitative analyses, they suggest that previous estrogen deficiency establishes a distinct cellular state that may contribute to the greater osteoclastogenic responsiveness of OVX-derived macrophages following coronavirus infection.

In agreement with the *in vivo* findings, infection increased RANK, TRAF6 and NFATc1 expression, together with TNF and CCL5 production, in macrophages from both Sham and OVX animals, supporting activation of osteoclastogenic and inflammatory pathways by MHV-A59. TNF is a well-established promoter of osteoclastogenesis and has also been implicated in coronavirus-associated bone resorption [20,66]. Moreover, the greater production of IL-6 and CCL3 by infected OVX-derived cells suggests that estrogen deficiency further enhances the inflammatory signals that associated with the osteoclastogenic response, consistent with the recognized contribution of IL-6 to estrogen deficiency-associated bone loss [67–69]. Importantly, the increased expression of cathepsin K only in infected OVX-derived cells further supports the acquisition of a more resorptive phenotype, as this protease is the major mediator of organic bone matrix degradation and is enhanced under estrogen-deficient conditions [57,70].

On the other hand, the response to exogenous RANKL further clarifies the nature of this effect. Although MHV-A59 infection did not trigger increase in TRAP^+^ numbers, OVX-derived cells stimulated with RANKL generated more osteoclasts than Sham-derived cells, and presented higher IL-6, CCL5 levels, and cathepsin K expression. The persistence of a higher inflammatory and cathepsin K response indicates that previous estrogen deficiency may influence the functional phenotype of osteoclasts even when differentiation is already strongly driven by RANKL. CCL3 and CCL5 may further reinforce this phenotype by promoting osteoclast precursor differentiation, and survival, while CCL5 can also participate in the coupling between bone resorption and formation [41–43,49,71–72]. Thus, together with the *in vivo* data, these results support a model in which MHV-A59 provides an acute osteoclastogenic stimulus that is amplified by the previous estrogen-deficient environment, involving RANKL-independent activation of osteoclastogenic and inflammatory pathways and resulting in a more resorptive cellular phenotype.

In conclusion, our findings demonstrate that MHV-A59 infection accelerates bone loss in ovariectomized mice during the acute phase of infection, indicating that estrogen deficiency transiently increases skeletal vulnerability to viral infection. *In vitro*, bone marrow-derived macrophages from OVX animals showed greater osteoclastogenic responsiveness to MHV-A59, even in the absence of exogenous RANKL, together with increased IL-6, CCL3, and cathepsin K, indicating that estrogen deficiency establishes a cellular state that amplifies the inflammatory and osteoclastogenic effects of infection. Although obtained in an experimental model, these findings may have clinical relevance, as COVID-19 often has worse prognosis in individuals with metabolic and systemic comorbidities, including diabetes, obesity, and renal dysfunction, which can independently impair bone remodeling and increase fracture risk [73–75]. Moreover, glucocorticoid therapy in severe cases may represent an additional skeletal risk by promoting bone resorption and suppressing bone formation [76–78]. Thus, the interaction between viral infection and pre-existing conditions that compromise bone homeostasis may contribute to a greater skeletal impact of COVID-19 than that observed in otherwise healthy individuals, highlighting the importance of considering viral infection as an additional modifier of bone remodeling and investigating its consequences in populations already susceptible to bone loss.

## DATA AVAILABILITY

The data with the findings of this study are available from the corresponding author upon reasonable request.

## ACKNOWLEDGMENTS

The authors thank Ilma Marçal de Souza, Letícia Soldati Silva, Rosemeire Oliveira, Tania Colina and Frankcinéia Assis for their technical support. The authors also acknowledge the support and the use of facility at the Centro de Microscopia da UFMG. This work was supported by the National Institute of Science and Technology in Dengue and Host-Microorganism Interactions (INCT Dengue e Interação Microrganismo-Hospedeiro; CNPq, Brazil; process 408527/2024-2; and FAPEMIG, Brazil; process APQ-08695-25), the Coordenação de Aperfeiçoamento de Pessoal de Nível Superior (CAPES, Brasil; Finance Code 001). Additional support was provided by FAPEMIG processes APQ-04650-23 and APQ-02618-23. Artificial intelligence (AI) tools were used to improve the English language, grammar, and clarity of the manuscript. The authors remain fully responsible for the content and scientific accuracy of the manuscript.

---

**Supplementary Figure 1. Effects of ovariectomy and MHV-A59 infection on clinical parameters and pulmonary alterations.** (A) Experimental design. (B) Percentage variation of body mass over time. (C) Variation in body mass in the first days after infection, evidencing the acute phase of the disease. (D) Leukocyte count and (E) granulocyte/lymphocyte (G/L) ratio throughout the course of infection (F) Uterine weight at 2, 5, and 60 days post-infection (dpi). (G) Pulmonary histopathological score of Sham and OVX mice inoculated with saline or infected with MHV-A59 at 2, 5, and 60 dpi. (H) Representative images of H&E-stained lung sections at 4× and 40× magnifications (5 dpi). Pulmonary levels of CXCL-1 (I), CCL2 (J), CCL3 (K), and CCL5 (L). White bars indicate control animals (saline) and blue bars indicate animals infected with MHV-A59. Points represent values for individual animals (n = 5), and bars indicate mean ± standard error of the mean (SEM). Statistical differences are indicated by * (*p* < 0.05 vs. Sham + Saline), # (*p* < 0.05 vs. Sham + MHV-A59) and & (*p* < 0.05 vs. OVX + Saline).

**Supplementary Figure 2. Cortical bone microarchitecture at 60 days post-infection.** (A) Bone area (B.Ar), (B) total cross-sectional area (T.Ar), (C) bone area fraction (B.Ar/T.Ar), (D) bone perimeter (B.Pm), (E) total perimeter (T.Pm), and (F) cortical thickness (Cs.Th). Representative cross-sectional micro-CT images of the femoral cortical bone (G). White bars indicate control animals (saline) and blue bars indicate animals infected with MHV-A59. Points represent individual animal values (n = 5), and bars indicate mean ± standard error of the mean (SEM). Statistical differences are indicated by * (*p* < 0.05 vs. Sham + Saline), # (*p* < 0.05 vs. Sham + MHV-A59) and & (*p* < 0.05 vs. OVX + Saline).

