## Supplementary figures and images for "CORONAVIRUS INFECTION ACCELERATES EXPERIMENTAL OSTEOPOROSIS THROUGH THE ACTIVATION OF OSTEOCLASTS"

### Figure S1

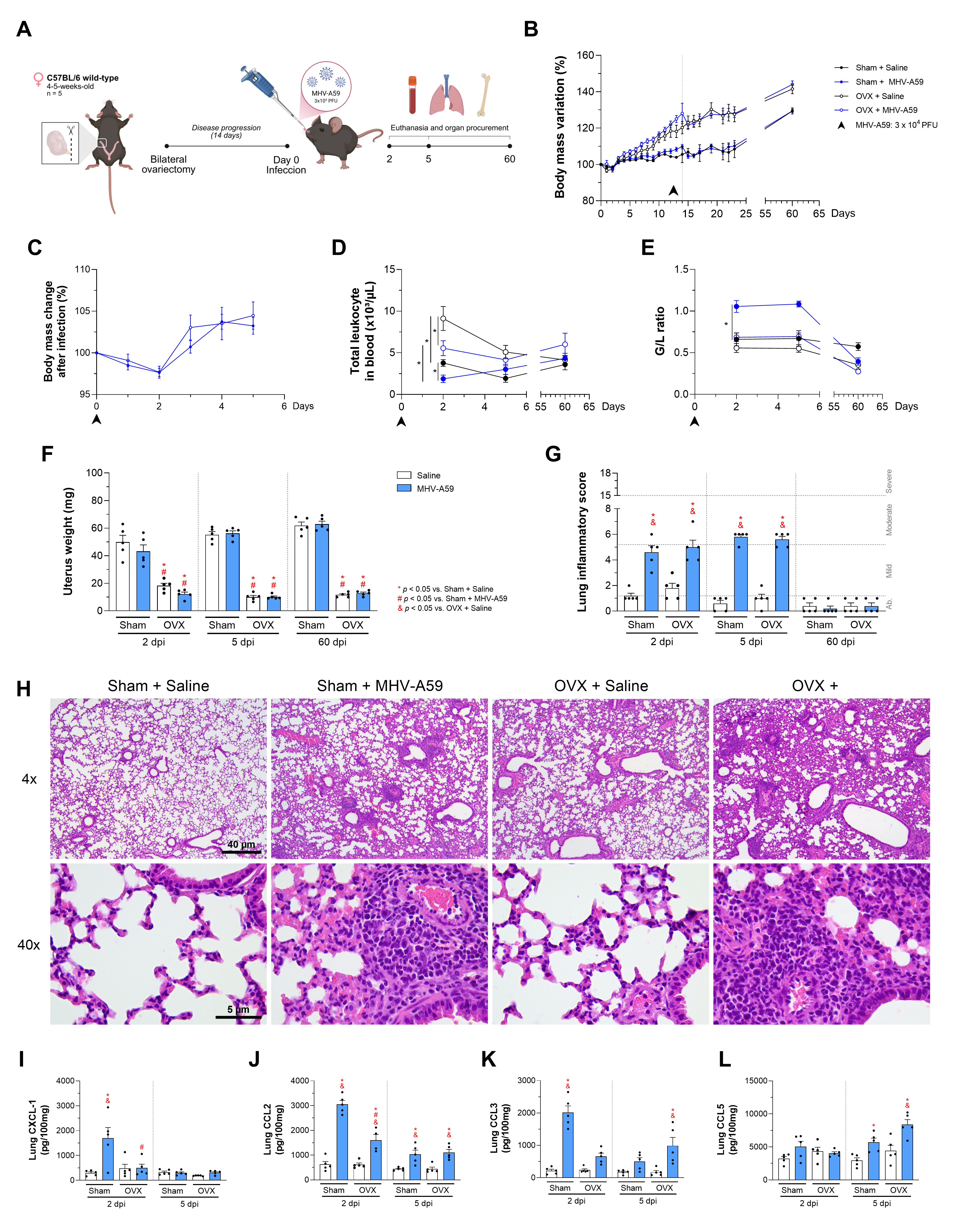

### Figure S2

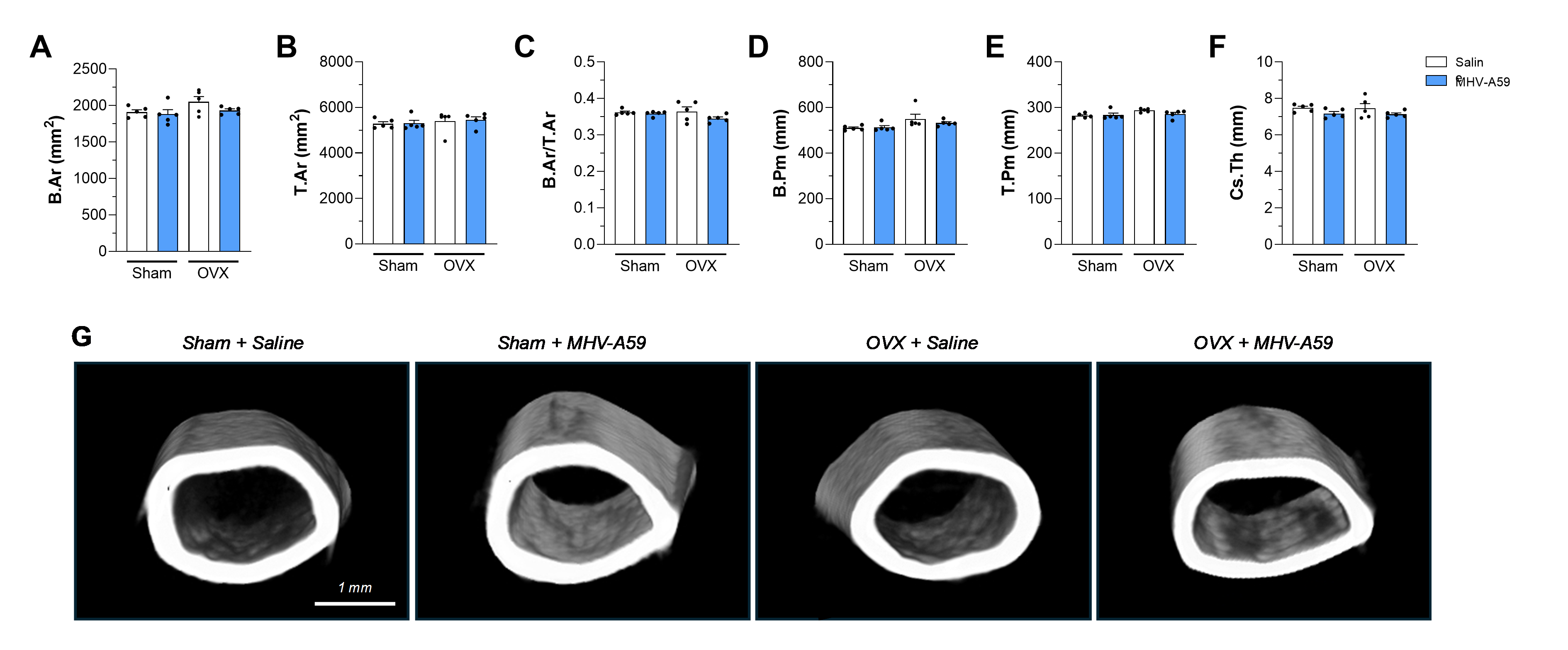
